# Translational profiling of microglia reveals artifacts of cell sorting

**DOI:** 10.1101/135566

**Authors:** Silvia S. Kang, Kelsey E. Baker, Xuewei Wang, Jeanne-Pierre Kocher, John D. Fryer

## Abstract

Microglia are the resident innate immune population of the central nervous system that constantly survey and influence their local environment. Transcriptomic profiling has led to significant advances in our understanding of microglia in several disease states, but tissue dissociation and purification of microglia is known to lead to cellular activation. Here we use RiboTag translational RNAseq profiling to demonstrate that commonly used cell sorting methods lead to a fundamental alteration of the microglial transcriptome, with several transcripts that can be used to mark artifacts of isolation. Microglial RiboTag RNAseq profiling after peripheral immune challenge with lipopolysaccharide demonstrates unique transcriptional targets that are not evident using cell sorting methodology. Finally, we applied our technique to reveal novel shared and distinct pathways when comparing microglial transcriptomes after peripheral challenge with bacterial or viral mimetics. This study has broad implications for approaches that examine microglial transcriptomes in normal and pathological states.

**Summary:** Kang et al. demonstrate artifactual induction of microglial transcripts associated with cell sorting. Using RiboTag translational profiling methodology, several markers of cell sorting artifact were revealed. Furthermore, RiboTag isolation unveiled changes in microglial transcriptomes following systemic inflammation that would otherwise have been masked by artifacts of cell sorting.

## Introduction

Microglia, the central nervous system (CNS) resident innate immune cell population, are critical cellular mediators during both normal homeostasis and disease pathogenesis (Davalos et al., 2005; Nayak et al., 2014; Nimmerjahn et al., 2005; Saijo and Glass, 2011). Under basal conditions, microglia comprise approximately 5 to 12% of the cellular population in different brain regions and are highly dynamic in nature allowing them to continually survey the CNS milieu and participate in synaptic pruning and shaping of neuronal circuits during homeostatic and inflammatory conditions (Hong et al., 2016a, 2016b; Lawson et al., 1990; Miyamoto et al., 2013, 2016; Nimmerjahn et al., 2005; Paolicelli et al., 2011; Schafer et al., 2012, 2012; Vasek et al., 2016; Zhan et al., 2014). They act as CNS sentinels and become rapidly activated during pathological states such as injury, inflammation, and neurodegeneration (Colonna and Butovsky, 2017; Klein et al., 2017; Nayak et al., 2014; Saijo and Glass, 2011). Microglial production of pro-inflammatory mediators such as tumor necrosis factor (TNFα), interleukin 1α (IL-1α), and C1q also induces neurotoxic A1 astrocytes, demonstrating there are also indirect microglial mechanisms for neuronal death (Liddelow et al., 2017). Additionally, peripheral viral or bacterial mimetic challenge can impact microglia and exacerbate neurodegeneration (Hoogland et al., 2015; Krstic et al., 2012; Perry et al., 2007) suggesting that even systemic inflammation shapes microglial function.

Due to the numerous roles of microglia in health and disease, it has become critical to illuminate microglial transcriptome signatures to elucidate important functional pathways. A microglial “sensome” that encompasses transcripts involved in endogenous ligand and microbial recognition has been defined using direct RNA sequencing and has been shown to distinctly change between adulthood and aging (Hickman et al., 2013). Additionally, signatures unique to microglia, relative to peripheral macrophages/monocytes, have been discovered by several groups to help define transcripts that allow for a clearer distinction of microglial versus infiltrating cellular responses (Bennett et al., 2016; Butovsky et al., 2014; Hickman et al., 2013). Isolation methodologies for microglial transcriptome studies invariably involve tissue dissociation and an enrichment process such as cell sorting or immunopanning. Recently, a novel Tmem119-based immunopanning method was developed to study microglial responses and will likely be an important tool to uncover microglial function in the future (Bennett et al., 2016). Importantly, there are currently no markers to determine if cell enrichment procedures alter microglial transcriptomes in order to increase the reliability of RNAseq datasets. In contrast to cell enrichment, a novel alternative approach to cellular transcript isolation is to use the RiboTag translational profiling system (Sanz et al., 2009). In this approach, Cre/Lox conditional expression of an epitope-tagged core ribosomal protein (RPL22) can be expressed in cell type of interest with the appropriate Cre recombinase line. For our study, we used the well-described tamoxifen inducible *Cx3cr1-Cre*^*ER*^ crossed to RiboTag mice to profile microglia. This allows for rapid extraction of microglial specific transcripts to circumvent issues associated with the cell enrichment or sorting such as enzymatic digestion and long preparation times. Additionally, this method allows for recovery of transcripts present in microglial processes that are likely lost during the process of creating single cell suspensions (Sanz et al., 2009).

Here we demonstrate that rapid isolation of transcripts from microglia using RiboTag prevents artifacts that may be introduced through long isolation protocols combined with cell sorting. Although both methods yield a common set of microglial specific transcripts, the cell enrichment and sorting protocol artificially upregulated numerous transcripts including those associated with systemic inflammation. Direct comparison of RNAseq datasets from sorted microglia with RiboTag microglia revealed robust and fundamental differences in transcriptomes. These differences were apparent both at baseline and after peripheral challenge with lipopolysaccharide (LPS). Several artifactual transcripts were identified that may be useful for investigators to assess whether their microglial enrichment via cell sorting or other purification methodologies may be inducing microglial activation. Furthermore, examination of RiboTag procured microglia transcriptomes demonstrated that systemic inflammation induced by systemic TLR3 or TLR4 challenge significantly altered microglial transcriptomes from baseline *in vivo*, with a broader induction of genes associated with TLR4 challenge. Our data highlight fundamental differences in interpretation of microglial transcriptomic data depending on the method of RNA isolation.

## Results and discussion

### Activation of microglia following TLR3 or TLR4 stimulated peripheral inflammation

Following systemic LPS challenge, there is a significant rise in pro-inflammatory cytokines in the CNS (Erickson and Banks, 2011; Kang et al., 2017). To determine when microglia become activated, C57BL/6J mice were challenged with 2μ g/g LPS intraperitoneally (i.p.). At 2, 4, 8, 24, 48, and 72 hours post injection, PBS perfused brains were isolated to assess transcript levels of *Il1b* and *Tnfa* proinflammatory cytokines and *Aif1* (encodes the microglial marker IBA1). Although there was an early rise in expression of both *Il1b* and *Tnfa* at 2 and 4 hours, only *Tnfa* was significantly elevated at the 24 hour time point (Fig. 1a,b). Notably, *Aif1* transcripts levels were not significantly increased until 24 hours post injection (Fig. 1c). To ensure that microglia are activated by TLR3 or TLR4 induced systemic inflammation, mice were challenged with either 2μ g/g LPS or 12μ g/g polyinosinic:polycytidylic acid (poly I:C) i.p. and examined for IBA1 expression by immunohistochemistry at 24 hours post injection. Microglia in both LPS and poly I:C conditions demonstrated an activated morphology and *Aif1* transcript levels were significantly increased over saline controls (Fig. 1d, e). The delayed *Aif1* induction relative to *Il1b* and *Tnfa* suggests that other cells, such as endothelial cells that sense LPS or poly I:C, may be mediating the early inflammatory response (Moser et al., 2016). Interestingly, activation of cells in the brain-immune interface, such as vascular endothelial cells, choroid plexus stromal and epithelial cells, as well as meningeal cells, have recently been shown to be early producers of chemokines and cytokines (e.g. CCL2, CXCL1, CXCL2 and IL-6) that are proposed to activate astrocytes (Hasegawa-Ishii et al., 2016). This indicates that indirect stimulation of glial cells via the brain immune interface may be a main mechanism for how peripheral inflammation mediates glial reactivity.

**Figure 1.**
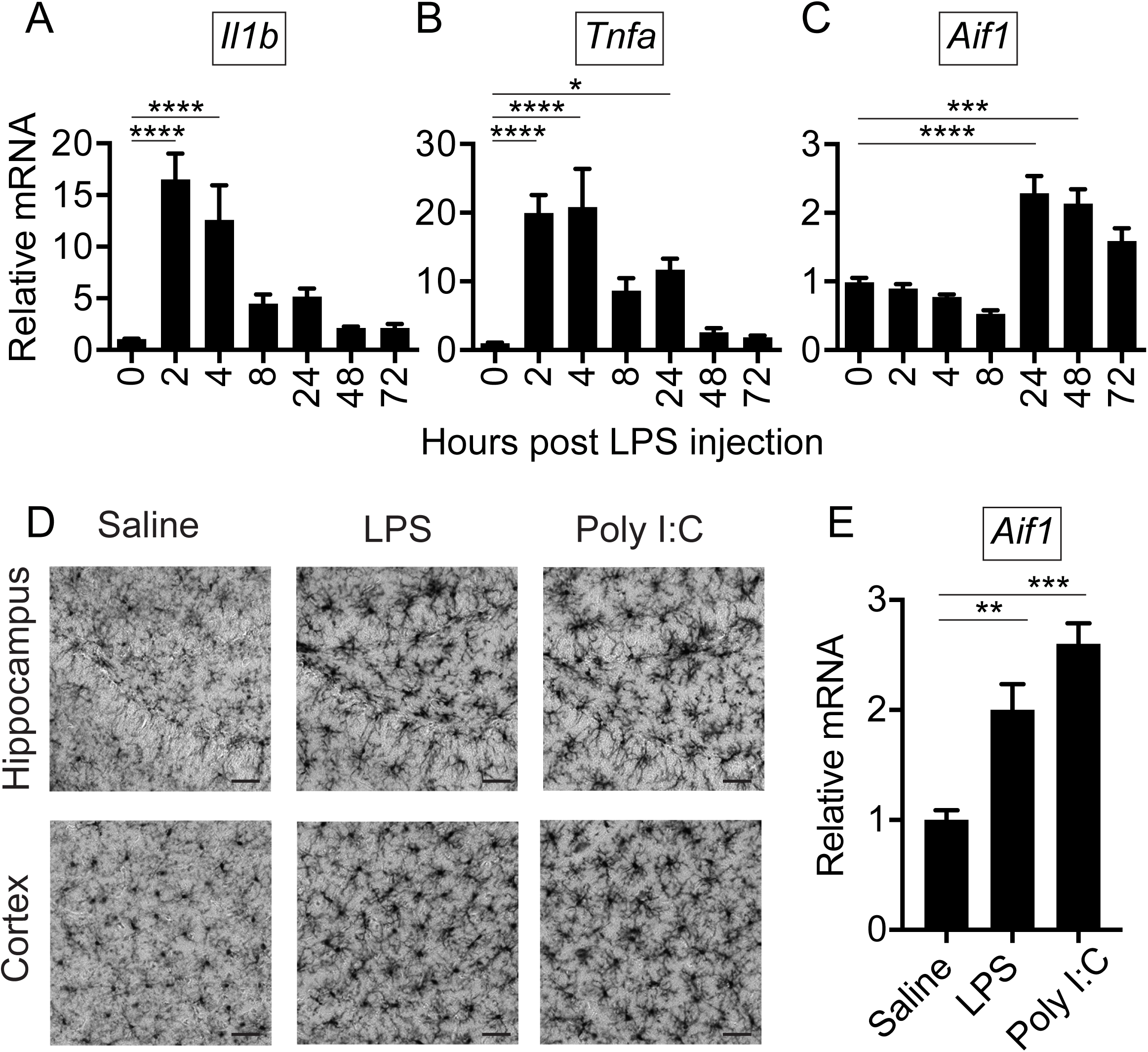
Systemic challenge results in microglial activation following the initial wave of pro-inflammatory cytokine production. C57BL/6J mice were injected with 2μ g/g LPS i.p. and PBS perfused brains were harvested at 2, 4, 8, 24, 48, and 72 hours post challenge. RT-qPCR for (A) *Il1b*, (B) *Tnfa*, and (C) *Aif1* transcript levels in hemi-brain tissue. Shown are the averages ± S.E.M. for N=3-4 animals. *<p0.05, ***p<0.001, and ****p<0.0001 by one-way ANOVA with Tukey posthoc t-tests. N=2 independent experiments were conducted. C57BL/6J mice were injected i.p. with either saline, 2μ g/g LPS, or 12μ g/g poly I:C and PBS perfused forebrains were harvested at 24 hours. (D) Representative immunohistochemistry for IBA1 expression in 50μ m coronal sections in the hippocampus and cortex regions are shown (scale bar = 40 microns). N=4 mice per group, 2 independent experiments. (E) RT-qPCR performed for *Aif1* (encodes IBA1) showed increased microglial activation. Shown are the averages ± S.E.M. for N=4 animals. **<p0.01, ***p<0.001, by one-way ANOVA with Tukey posthoc t-tests. N=2 independent experiments were conducted.

### Microglia isolated using RiboTag technology reveals artifacts in the microglial transcriptome due to cell sorting

In order to determine if microglial transcripts are altered based on the RNA isolation procedure, we isolated RNA using both cell enrichment/cell sorting and a novel RiboTag approach. RiboTag Rpl22 mice express loxP sites flanking the endogenous terminal exon 4 of the Rpl22 gene (a ubiquitous core ribosomal subunit) that can be removed through Cre mediated recombination allowing for transcription of a repeated exon 4 of Rpl22 but engineered to contain a hemagluttanin (HA) epitope tag on the C-terminus (Sanz et al., 2009). We utilized the well characterized microglial mouse line with a tamoxifen inducible Cre and eYFP, *Cx3cr1-Cre*^*ERT*2^*-IRES-eYFP,* under the control of the *Cx3cr1* promoter (Parkhurst et al., 2013), referred to as Cx3cr1^CreER^. To ensure that HA expression was restricted to microglia, double heterozygous Cx3cr1^CreER/+^; Rpl22^HA/+^ mice (referred to as RiboTag) were injected with 100μ g/g tamoxifen i.p. once daily for 3 consecutive days, to induce Cre recombinase mediated Rpl22-HA expression and PBS perfused brains were examined one week later by immunofluorescence. Confocal microscopy revealed perfect co-localization of *e*YFP encoded by the *Cx3cr1-CreER-IRES-eYFP* with HA epitope expression as well as with P2RY12 staining, confirming microglial expression of HA (Fig. 2A).

**Figure 2.**
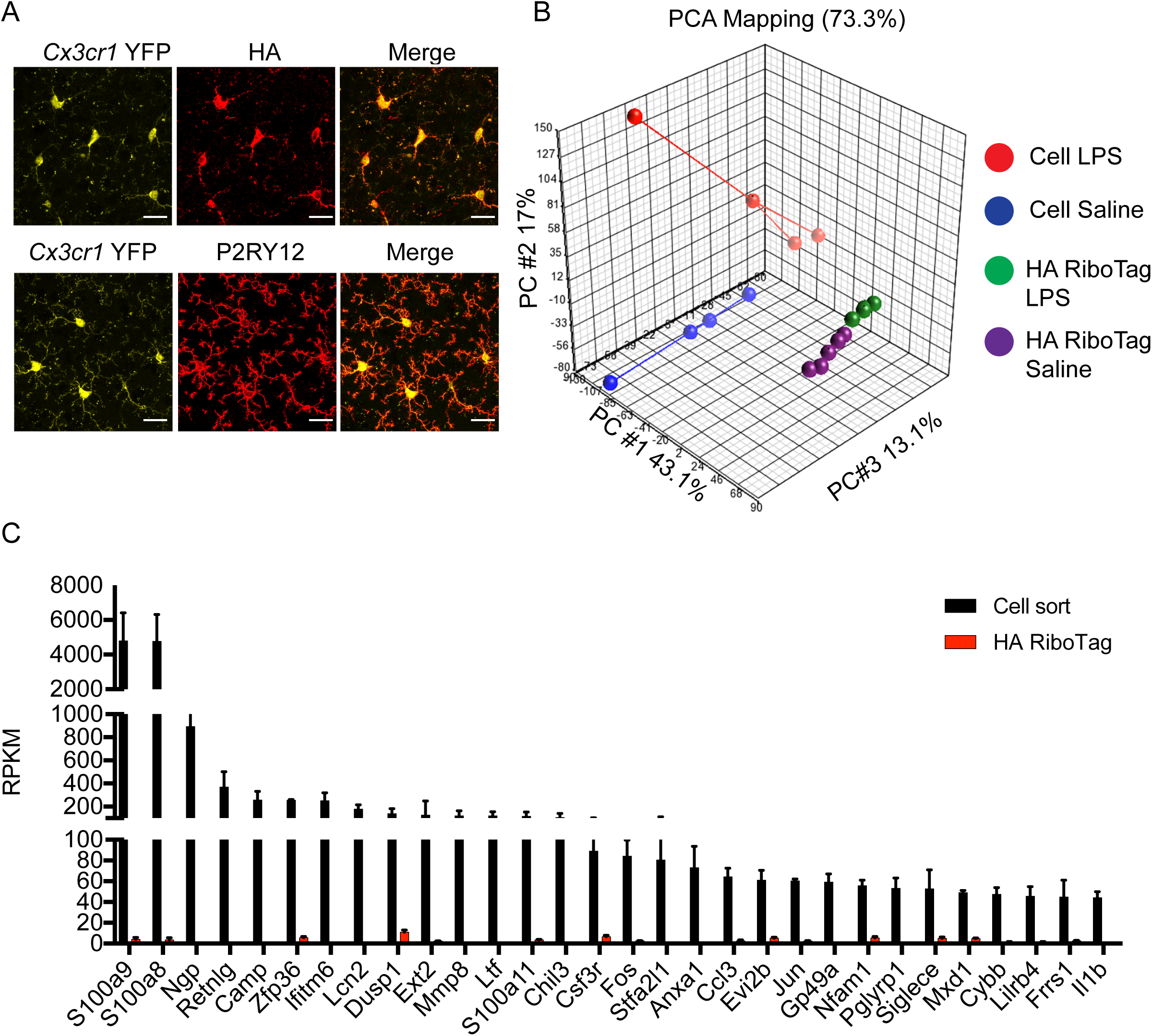
Microglial transcriptomes derived from RiboTag versus cell sorting reveal artifacts of cellular isolation. Cx3Cr1^CreER/+^; Rpl22^HA/+^ mice were injected with 100μ g/g tamoxifen i.p. once daily for 3 consecutive days to induce Cre recombinase mediated Rpl22^HA^ expression. (A) Confocal analysis of 50μ m coronal sections stained with anti-HA or anti-P2YR12 antibodies revealed co-localization of EYFP driven by the endogenous *Cx3cr1* promoter along with both HA and P2YR12 staining. A representative image is shown from N=3 experiments (scale bar = 20 micron). (B) Principal component analysis (PCA) of RNAseq datasets of microglial transcriptomes from Cx3Cr1^CreER/+^; Rpl22^HA/+^ tamoxifen treated mice isolated using either the RiboTag method or by cell sorting for CD11b^+^CD45^lo/int^ microglia at 24 hours after injection with either saline or 2μ g/g LPS. Centroids for each group are indicated on the PCA plot. N=3-4 animals per group. (C) The top 30 most abundant microglial transcripts enriched in cell sorted microglia (> 10 fold enriched over input) that are also over 10 fold higher RPKM relative to RiboTag isolated transcripts demonstrates the magnitude of cellular artifacts for the top 30 transcripts. Shown are the averages ± S.E.M. from the N=3-4 animals per group. Data analyzed by t-test with two-stage linear step-up procedure of Benjamini, Krieger and Yekutieli with false discovery rate (FDR) set at 5%.

Due to the rapid nature of RiboTag mediated isolation, and the ability to obtain transcripts from even fine processes that would likely be lost during isolation from cell sorting, we next compared data yielded from both these methods. Tamoxifen treated RiboTag mice were injected with either saline or 2μ g/g LPS i.p. and PBS perfused brains were either used to isolate microglia using the RiboTag method or isolated via cell enrichment combined with isolation by flow cytometric cellular sorting. For the RiboTag method, mice were PBS perfused and the tissue was immediately disrupted with sonication in a buffer containing cyclohexamide to lock nascent mRNA onto ribosomes and RNasin to prevent RNA degradation, effectively instantaneously crosslinking ribosomes to RNA(Doyle et al., 2008; Sanz et al., 2009). Isolation of microglial specific transcripts was conducted by incubation of transcripts with an anti-HA antibody followed by magnetic bead separation from non-HA tagged ribosomes. For cell sorting, PBS perfused brain tissue was enzymatically digested and microglia were enriched from a 60/40 percoll gradient to remove myelin debris (Kim et al., 2009). Cells were stained with CD11b and CD45 and sorted for the microglial CD11b^+^CD45^lo/int^ population to >98%. In opposition to the immediate removal of RNA transcripts from the cellular source that occurred within seconds of terminal perfusion using the RiboTag methodology, cell enrichment/cell sorting required hours of isolation prior to cell lysis. Microglial transcriptomes were compared between four groups (n=3/group): RiboTag-saline, RiboTag-LPS, sorted-saline, sorted-LPS. RNAseq transcriptomes were analyzed using a multiple t-test, two-stage step-up Benjamini-Krieger-Yekutieli method with an FDR set at 5% and q<0.05 considered significant.

Principle component analysis (PCA) demonstrated distinct populations that were separated not only by LPS treatment but also cell sorting versus RiboTag isolation in the saline treated groups (Fig. 2B). To examine microglial associated genes, the abundance of transcripts derived from cell sorting or RiboTag were compared to total whole brain input values from the same samples. Transcripts that were >10 fold over input were considered to be microglial enriched by either method. Interestingly, the top 25 most abundant microglial transcripts determined by cell sorting revealed numerous transcripts that were either not significantly enriched in microglia or missed the >10 fold cut off criteria in the RiboTag dataset. Conversely, almost all of the top 25 most abundant transcripts enriched in microglia from the RiboTag dataset were also enriched in transcriptomes from cell sorted microglia, with the one exception of *Sepp1* which was enriched 9.98 fold and just barely missed the >10 fold cut off (Table I, Supplemental Table I). These data demonstrate that for the most abundantly expressed microglial genes, RiboTag methodology identified transcripts that are also identified by cell sorting. However, cell sorting identified numerous transcripts that were not identified with RiboTag isolation and are likely an artifact of cellular isolation. The RiboTag method was further validated by the fact that many of the most abundant transcripts found using this method (e.g. *Hexb, P2ry12, Csf1r, Tmem119,* and *Ctss*) have previously been identified as microglial associated transcripts and proteins (Bennett et al., 2016; Butovsky et al., 2014; Hayashi et al., 2013; Hickman et al., 2013).

**Table I:**
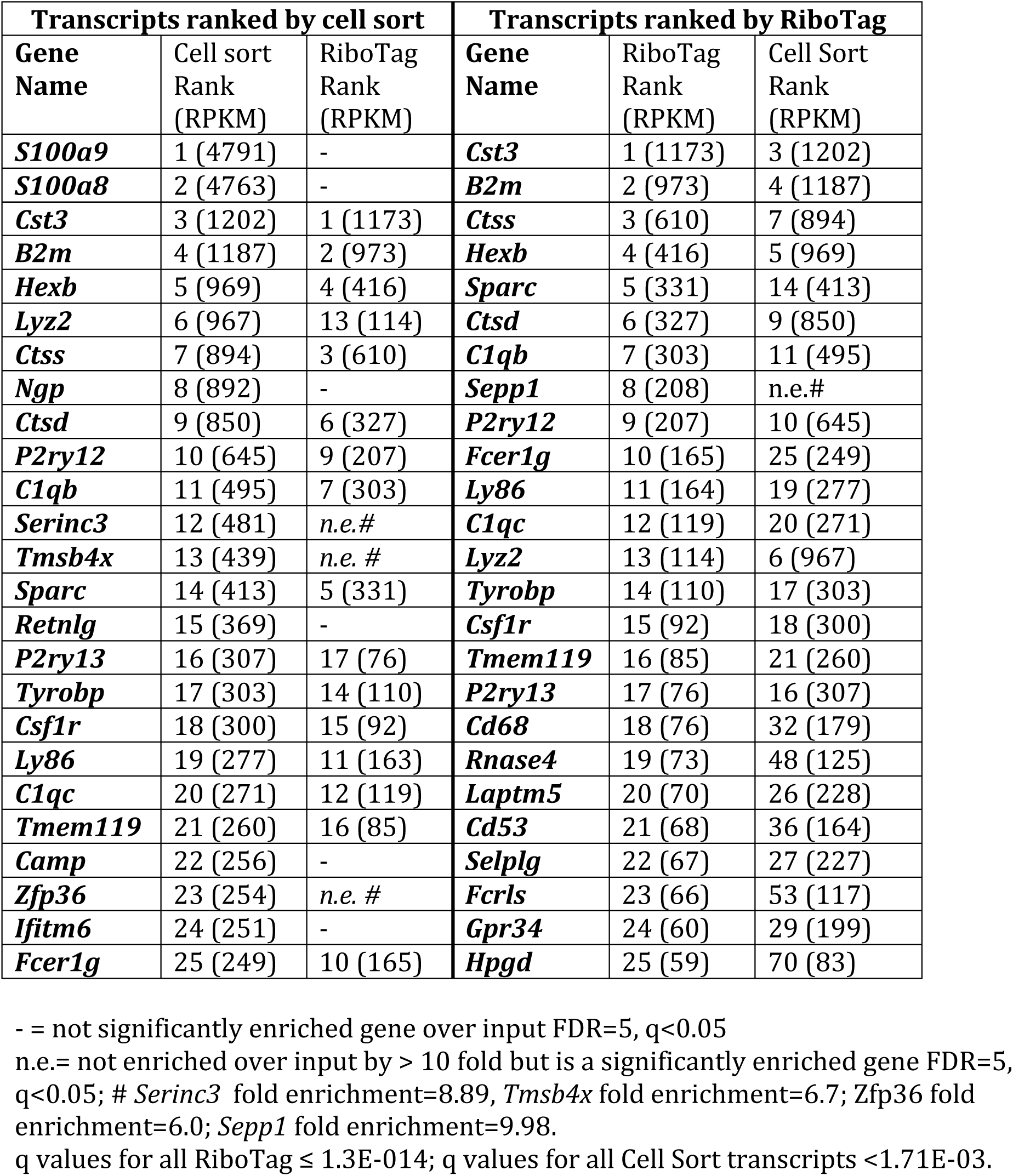
Comparison of the top 25 transcripts enriched in microglia from cell sort dataset versus RiboTag dataset (enrichment defined as RPKM>10 fold over input RPKM), ranked by absolute abundance (RPKM).

The PCA analysis demonstrated a clear segregation between cell sorted versus RiboTag isolated microglial transcriptomes. Examination of significantly enriched microglial transcripts identified by cell sorting but which are expressed an additional 10 fold higher in overall abundance RiboTag datasets (i.e. the most artifactually upregulated genes from cell sorting), yielded 297 transcripts. Importantly, 73 of these transcripts were not significantly enriched with RiboTag isolation at all such as *S100a8*, *S100a9*, *Ngp*, *Retnlg*, *Ifitm6*, *Lcn2*, *Dusp1*, *Mmp8*, *Ltf*, *Chil3*, *Fos*, and *Jun*, suggesting that these may be critical markers that may be useful in determining whether isolation methodologies are aberrantly activating microglia (Fig. 2C, Supplemental Table I). Previously, an invaluable database for examination of transcripts based on CNS cell type was published along with a searchable website (Zhang et al., 2014). With the exception of *Lcn2*, *Jun*, *Fos*, and *Dusp 1*, which were also expressed in additional cell types, the remaining transcripts above appeared to be predominantly expressed by microglia using a panning method to purify this population. Interestingly, a recently released updated dataset of the microglial transcriptome, based on a new protocol utilizing a Tmem119 specific antibody for microglial purification by cell sorting, no longer appears to have increased expression of *S100a8*, *S100a9*, *Ngp*, *Retnlg*, *Ifitm6*, *Lcn2*, *Dusp1*, *Mmp8*, and *Ltf* (Bennett et al., 2016). These data are in accordance with our RiboTag results and are supportive of the potential of transcripts such as *S100a8*, *S100a9*, *Ngp*, *Retnlg*, *Ifitm6*, *Lcn2*, *Dusp1*, *Mmp8*, and *Ltf* to act as indicators of potential artifacts induced by microglial isolation. Increased expression of *S100a8* and *S100a9*, also known as *Mrp8* and *Mrp14* respectively, have been shown during phagocyte activation and they are associated with the induction of a stress tolerance state (Austermann et al., 2014; Vogl et al., 2007). Within the CNS, *S100a8* and *S100a9* have been associated with CD11b^+^ cells surrounding amyloid plaques and Mmp8 is detected in microglia during neuroinflammation (Han et al., 2016; Kummer et al., 2012). This indicates that these transcripts are likely induced during an activation state, and in this case, are also induced by the cellular isolation process. Additional markers, such as the transcription factors *Fos* and *Jun*, appear to be enriched in microglia over whole brain lysates in the updated Tmem119-based dataset, potentially reflecting early markers of a shifting transcriptome due to cellular isolation. Interestingly, these transcripts were not enriched in our RiboTag dataset and were found to be either equally and more highly expressed in endothelial and astrocytes, respectively, relative to microglia in the published database (Zhang et al., 2014), suggesting expression was likely based on activation rather than serving as a microglia specific transcript. Notably, the new Tmem119-based microglial isolation methodology, which lacks many of the artifactual transcripts we defined here, involves cell sorting demonstrating that cell sorting in it of itself may not induce artifacts, rather the entire enrichment protocol needs to be taken into account.

Seminal work defining a microglial sensome and molecular signatures have been conducted using deep sequencing to define the microglial transcriptomes (Butovsky et al., 2014; Hickman et al., 2013). Interestingly, several transcripts that emerged as some of the most abundant microglia associated transcripts, including *Cst3*, *B2m*, *Ctss*, *Ctsd*, and *C1qb* were not observed in the initial characterization of the sensome. These are also transcripts that were highly expressed in the Tmem119-based isolation dataset (Bennett et al., 2016). The basis for these differences is unclear, but it suggests that different isolation methodologies may be causing significant differences in the microglial transcriptomes, making a marker for artifact introduction by methodology even more pressing.

### Cell sorting masks transcriptome activation in microglia after peripheral immune stimulation with LPS

We next asked the question whether our RiboTag methodology would reveal differences in transcriptome after peripheral LPS challenge compared to cell sorting. Our analysis of microglial transcripts that were derived from cell sorted vs RiboTag purified methods after LPS challenge revealed a striking discrepancy between the responses to systemic inflammation based on isolation method. PCA revealed distinct clustering of cell sorted microglia after LPS compared to RiboTag microglia after LPS treatment (Fig. 2A). Additionally, examination of the top 25 LPS induced transcripts, ranked by fold induction, revealed very little overlap between cell sorted vs RiboTag isolated transcripts (Fig. 3A,B, Supplemental Table II). For transcripts such as *Prok2* and *Cst7*, there appeared to be a “two-hit” effect where cell sorting alone had low to minimal impact on expression; however, in combination with microglial activation induced by peripheral LPS challenge, an aberrant response, *Prok2*, or a highly exaggerated response, *Cst7*, were revealed (Fig 3C,D). Additionally, while some transcripts were artifactually increased, others were dampened due to a higher baseline expression of the transcript caused by cell sorting. This was observed for both *Lcn2* and *IL1b*, which are known to be activated in the CNS with LPS challenge (Kang et al., 2017), where RiboTag isolated transcripts revealed a more robust LPS response relative to those that were masked in cell sorting derived transcripts (Fig. 3 E,F). In fact, the top 10 hits from the LPS cell sort dataset only has a single transcript in common with the top 10 hits from the LPS RiboTag dataset: *Saa3*. Thus, the experimental candidates that one might pursue in terms of mediating the biological effect (LPS in this case) would be fundamentally different depending on the methodology and could lead down the wrong path altogether.

**Figure 3.**
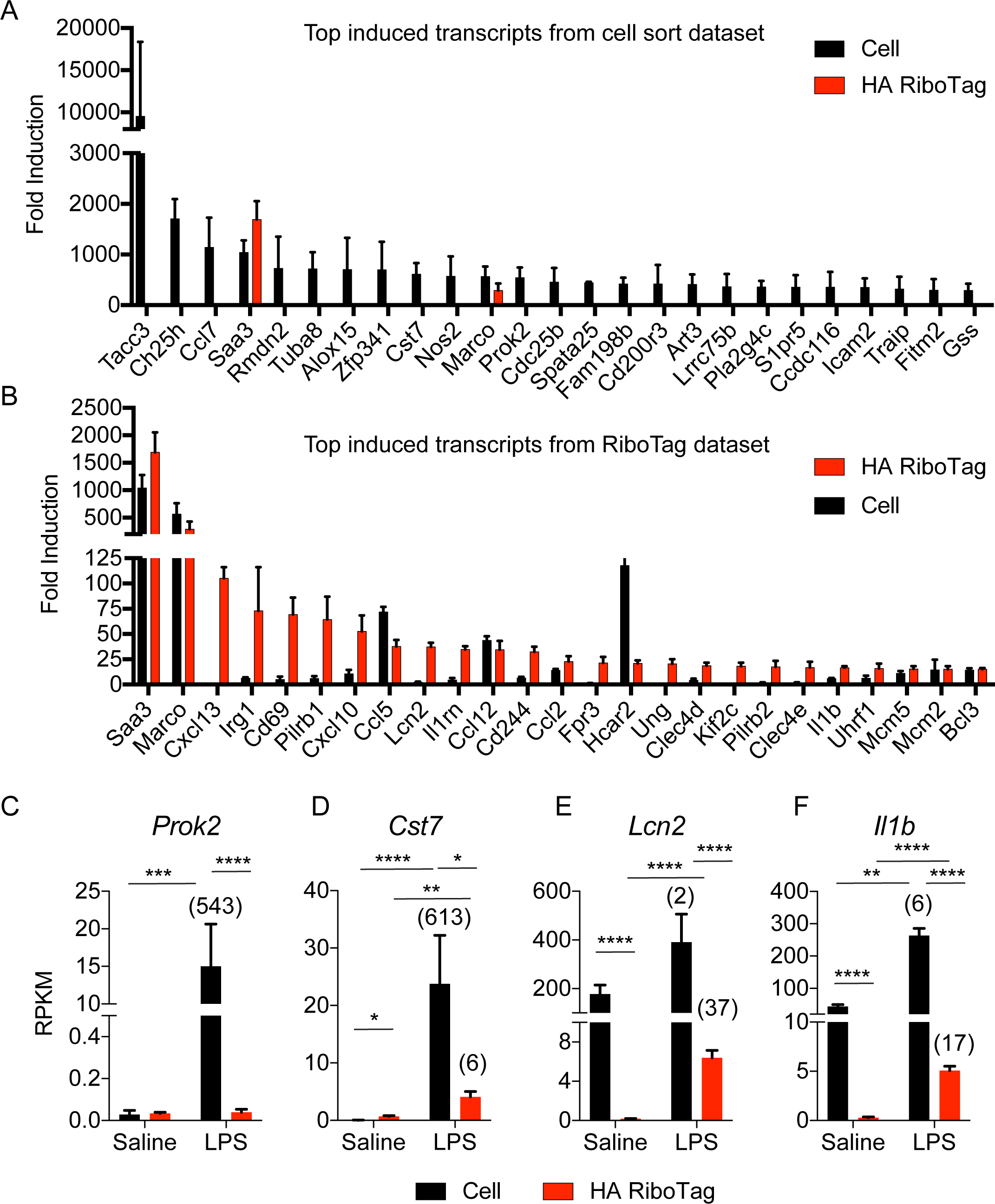
Artifacts from cell sorting alters the ranking of top microglial transcripts induced by systemic LPS stimulation. Cx3Cr1^CreER/+^; Rpl22^HA/+^ mice were injected with 100μ g/g tamoxifen i.p. once daily for 3 consecutive days to induce Cre recombinase mediated Rpl22 HA expression. One week later, animals were injected with 2 μ g/g LPS i.p. and harvested 24 hours later. Transcriptomes were isolated either using HA RiboTag or by cell sort method (cell sorted microglia came from identical Cx3Cr1^CreER/+^; Rpl22^HA/+^ mice injected with tamoxifen). The top 25 transcripts that were significantly induced by LPS ranked according to (A) fold induction in cell sorted microglia dataset or (B) fold induction in RiboTag isolated microglia dataset. Shown are the averages ± S.E.M. from N=3-4 animals per group. Data analyzed by t-test with two-stage linear step-up procedure of Benjamini, Krieger and Yekutieli with false discovery rate (FDR) set at 5%. Several transcripts were altered by cell sorting that demonstrate a “two hit” increase in expression after LPS, including (C) *Prok2*, (D) *Cst7*, or inductions that were masked due to increased basal levels by cell sorting, including (E) *Lcn2* and (F) *Il1b*. Shown are the averages ± S.E.M. from the N=3-4 animals per group, two-way ANOVA with Tukeys posthoc t-tests. Numbers in parentheses indicate fold induction compared to saline treated mice.

### Peripheral inflammation causes significant changes in the microglial transcriptome

Because peripheral inflammation can have lasting effects on CNS function, potentially through altered microglial function, it is important to understand how systemic inflammation shapes microglia in general (Hoogland et al., 2015; Krstic et al., 2012; Perry et al., 2007). To further test the utility of our RiboTag method, we profiled the microglial transcriptome at 24 hours following peripheral LPS (TLR4 agonist) or poly I:C (TLR3 agonist) induced systemic inflammation. A large change in the microglial transcriptome was observed after challenge with both LPS (1417 altered transcripts) and Poly I:C (1049 altered transcripts) suggesting that LPS induced a much broader response (Fig. 4A). Pathway analysis of significant genes induced by LPS or poly I:C challenge relative to saline controls revealed similar, and expected, immune function related pathway activation including innate immune response, immune system process, and inflammatory process pathways (Fig. 4b). However, despite having 664 overlapping transcripts in microglia following these inflammatory challenges (Fig. 4a), the overall response was quite different. Examination of the top 25 transcripts that were highly induced following LPS challenge demonstrated that many of these transcripts (e.g. *Marco*, *Cxcl13*, *Irg1*, *Cd69*, *Il1rn*, *Clec4e*, *Clec4d, Il1b*) were not observed with poly I:C challenge (Fig. 4c). Additionally, examination of the top 25 transcripts induced by poly I:C challenge revealed that although LPS challenge also induced several of these genes (e.g. *Ccl2*, *Ccl12*, *Cxcl10*, *Oas3*), the magnitude of the induction was much lower (Fig. 4d). A previous study examining microglial transcriptome responses to TLR3 or TLR4 stimulation demonstrated an induction of several immune response genes, including *Oas2*, *Oas3*, *Ifi35*, *Ifi203*, *Ifi204*, *Ifit1*, and *Ifit2* in both LPS and polyI:C challenge (Das et al., 2015; Diamond and Farzan, 2013; Samuel, 2001). In accordance with these results, we found these genes to be induced in both conditions, with the exception of *Oas2* and *Ifi203*, which were only observed in the poly I:C condition. Our data in Fig.1A demonstrates that inflammation precedes microglial *Aif1* transcript upregulation, suggesting that microglia may be indirectly activated by peripheral inflammation as was observed in astrocytes (Hasegawa-Ishii et al., 2016). Overall, these data indicate that microglial transcriptomes are significantly shaped by peripheral inflammation *in vivo*, with LPS inducing a much greater shift in gene expression compared to poly I:C.

**Figure 4.**
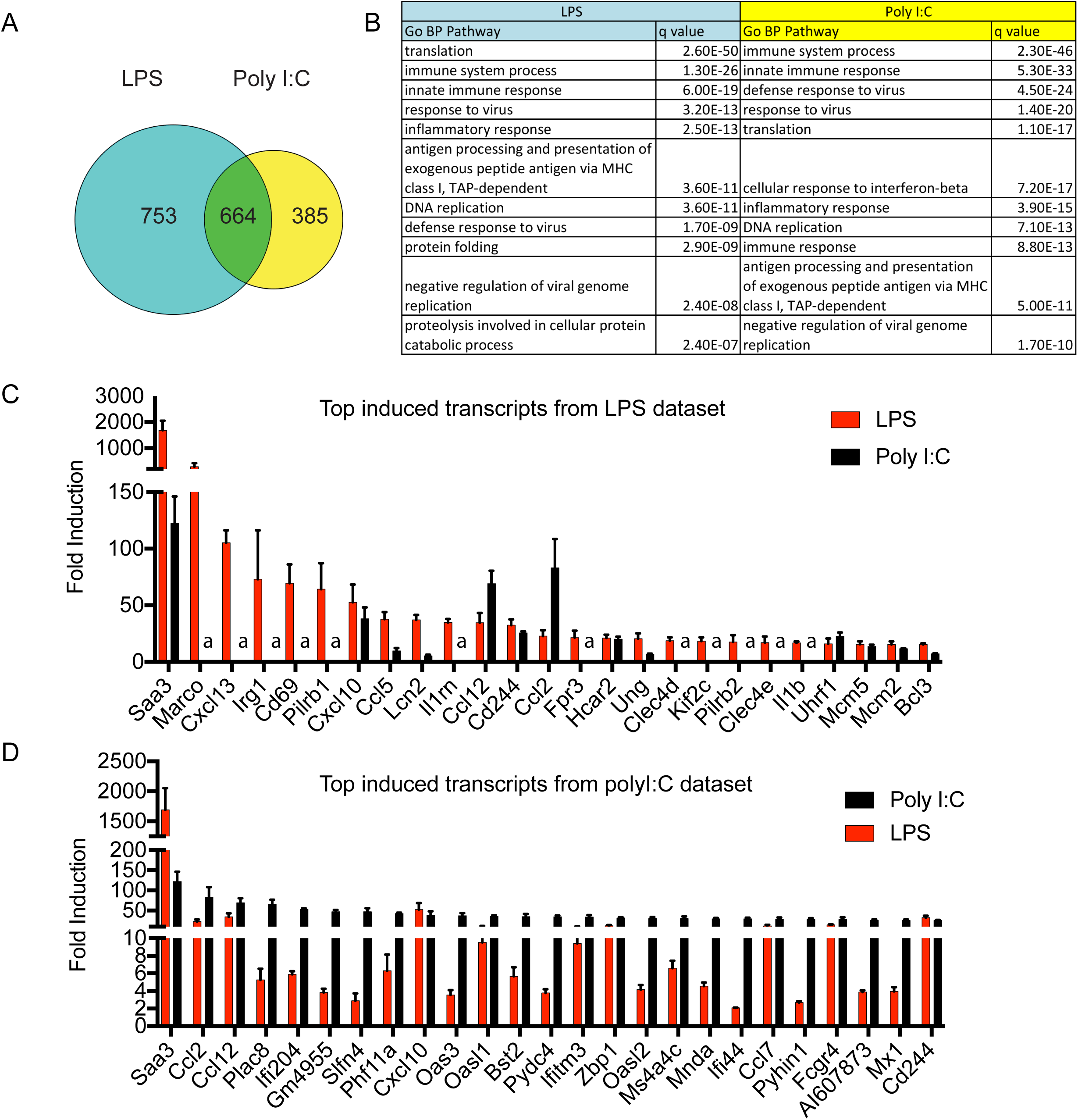
Microglial transcriptomes are differentially altered by peripheral inflammation induced by bacterial or viral mimetics. Cx3Cr1^CreER/+^; Rpl22^HA/+^ mice were injected with 100μ g/g tamoxifen i.p. once daily for 3 consecutive days to induce Cre recombinase mediated Rpl22 HA expression. One week after the last dose of tamoxifen, mice were injected i.p. with saline, 2 μ g/g LPS, or 12 μ g/g of Poly I:C and harvested 24 hours later. RNAseq analysis of microglial transcriptomes that were significantly altered by stimulation. Data analyzed by t-test with two-stage linear step-up procedure of Benjamini, Krieger and Yekutieli with false discovery rate (FDR) set at 5%. (A) Venn diagram demonstrating overlap of transcripts induced by both LPS and poly I:C. (B) Gene ontology biological process (GO BP) pathway analysis demonstrating distinct pathways enriched in microglia after LPS or poly I:C. The top 25 transcripts that were significantly induced in RiboTag datasets ranked according to (C) fold induction by LPS treatment or (D) fold induction by poly I:C treatment. a= transcripts that failed to pass the initial cutoff for log2 RPKM expression >-2.

In summary, our RiboTag method of profiling microglia demonstrates that cell sorting of microglia results in a fundamental alteration of their transcriptome both at baseline and after peripheral challenge with LPS. Several novel transcripts, including *S100a8*, *S100a9*, *Ngp,* etc may serve as important indicators of whether microglial isolation methodologies are introducing artifacts during purification. This provides critical target transcripts for investigators to use to identify if their own isolation procedures are likely to introduce artifact and will, as a result, make the resulting transcriptomic data more interpretable and reproducible. We have demonstrated that RiboTag profiling of microglia is an attractive alternative method to circumvent these issues, and have utilized this methodology to demonstrate that microglia have significant transcriptomic changes in response to peripheral inflammatory challenges. It is unknown whether profiling of cell sorted microglia in other conditions (e.g. aging, neurodegenerative disease, traumatic brain injury, infection, etc) has also led to the prioritization of the wrong targets, but utilization of RiboTag or other translational profiling methodologies will be necessary to definitively address this.

### Materials and Methods

#### Animals

C57BL/6J and *Cx3Cr1*^creER/+^; Rpl22^HA/+^ mice (The Jackson Laboratory, Bar Harbor, ME) were housed under standard laboratory conditions in ventilated cages on 12 hour light:dark cycles in a specific pathogen-free environment. To generate the *Cx3Cr1*creER; Rpl22 mice, two independent strains both on a C57BL/6J background derived from the Jackson Laboratory, the Cx3Cr1^tm2.1(Cre/ERT2)^ and Rpl22^tm1.1PSam^ were bred and used as heterozygous mice for all studies (i.e. *Cx3Cr1*^creER/+^; Rpl22^HA/+^). For LPS and poly I:C studies, animals were injected with 2μ g/g LPS (0111:B4; Sigma, Saint Louis, MO) or 12μ g/g poly I:C (HMW; Invivogen, San Diego, CA) intraperitoneally (i.p.). Animal protocols were reviewed and approved by Mayo Clinic Institutional Animal Care and Use Committee.

#### Tissue processing and RT-qPCR

Animals were deeply anesthetized with pentobarbital prior to cardiac perfusion with phosphate-buffered saline (PBS) to expunge blood from the cerebrovasculature. Forebrain tissues were quickly frozen on dry ice and stored at - 80°C until further processing. Tissues were briefly sonicated in Tris buffered saline with EDTA (TBSE) (50mM Tris pH=7.5, 150mM NaCl, 1mM EDTA) with 1x protease and phosphatase inhibitors (Thermo Scientific, Waltham, MA). An aliquot of the tissue suspension was processed for RNA using RNeasy Plus Mini Kit (Qiagen, Valencia, CA). Total RNA was isolated from sonicated tissues using an RNeasy Plus mini isolation kit according to manufacturer’s instructions. Random-primed reverse transcription was performed according to manufacturer protocols (Invitrogen-Life Technologies, Grand Island, NY). cDNA was added to a reaction mix (10μ L final volume) containing 300nM gene-specific primers and Universal SYBR green supermix (Biorad, Hercules, CA). All samples were run in triplicate and were analyzed on a Quant Studio 7 Flex Real Time PCR instrument (Applied Biosystems - Life Technologies). Relative gene expression was normalized to GAPDH controls and assessed using the 2^-ΔΔCT^ method. Primer sequences are as follows (5’ to 3’): GAPDH F: CTGCACCACCAACTGCTTAG, GAPDH R: ACAGTCTTCTGGGTGGCA GT, Aif1(Iba1) F: GGATTTGCAGGGAGGAAAAG, Aif1(Iba1) R: TGGGATCATCGAGGAATTG, Il1b F:CCTGCAGCTGGAGAGTGTGGAT. IL1b R: TGTGCTCTGCTTGTGAGGTGCT, Tnfa F:AGCCCACGTCGTAGCAAACCAC, Tnfa R: AGGTACAACCCATCGGCTGGCA.

#### Immunohistochemistry

PBS perfused hemi-brains were drop fixed into 10% neutral buffered formalin (Fisher Scientific, Waltham, MA) overnight at 4°C. Tissue was then placed in 30% sucrose (Sigma, Saint Louis, MO) dissolved in PBS overnight at 4°C. 50μ coronal brain sections were cut on a freezing-sliding microtome and stored in cryoprotectant at -20°C until staining. Sections were blocked for endogenous peroxidase activity and permeabilized with 0.6% H_2_0_2_, 0.1% NaN_3_ in PBS-X (1X PBS containing 0.3% Triton-X) for 30 minutes at RT. Sections were blocked with 1% milk in PBS-X followed by incubation with rabbit anti-IBA1 at 1:8000 (cat# 019-9741, Wako, Richmond, VA) in 0.5% milk PBS-X for 2 days at 4°C. Sections were then incubated with the Vectastain kit anti-rabbit IgG (Vector Labs, Burlingame, CA) overnight at 4°C followed by ABC component for 4hrs and developed using the DAB kit (Vector Labs) according to manufacturer’s instructions. Images were acquired using an Aperio XT Scanner (Aperio, Vista, CA) at a 20x magnification.

#### Cell sorting of microglia

PBS perfused brains from mice challenged with either saline or 2μ g/g LPS i.p. were incubated with collagenase D for 20 minutes at 37°C. Brains were immediately disrupted into a single cell suspension, filtered through a 100μ m filter, and washed with HBSS. Samples were centrifuged for 5 minutes at 1200 rpm. Supernatants were aspirated and the cell pellet was resuspended in 90% percoll/1x HBSS. A percoll gradient was made with additional layers of 60% percoll/1x HBSS, 40% percoll/1x HBSS, and HBSS. Gradients were centrifuged for 18 minutes at 1700 rpm with no brake and cells were isolated from the 60/40 interface. Cells were washed twice with HBSS, incubated with Fc block for 10 minutes on ice in 1% BSA/HBSS buffer, and then stained for 30 minutes with anti-CD11b and anti-CD45 (Biolegend, San Diego, CA). Cells were washed, sorted using CD11b^+^ CD45^lo/int^ as the gate to identify microglia, and sorted directly into a 1.5mL microcentrifuge tube. RNA was extracted using a Qiagen RNeasy Microkit. This process from brain extraction to sorting into a microcentrifuge tube takes approximately 4 hours.

#### RiboTag Microglial RNA isolation

*Cx3Cr1*^cre/+^; Rpl22^HA/+^ mice were injected for 3 consecutive days with 100μ g/g tamoxifen (Sigma, Saint Louis, MO) resuspended in corn oil to induce Cre and allow for HA tagged Rpl22 expression. One week following the final injection, animals were challenged i.p. either with saline, 2μ g/g LPS, or 12μ g/g poly I:C. At 24 hours post injection, animals were anesthetized with pentobarbital and PBS perfused prior to forebrain harvest and immediate tissue sonication in homogenization buffer (50mM Tris pH 7.4, 100mM KCl, 12mM MgCl2, 1% NP-40) supplemented with 1mM DTT, 1x protease/phosphatase inhibitors (Thermo Scientific), 200U/mL RNAsin, 100 μ g/mL cycloheximide, and 1mg/mL heparin. The cycloheximide serves to lock mRNA onto ribosomal complexes. Lysates were centrifuged at 4°C at 10,000 rpm for 10 minutes. Supernatants were incubated with anti-HA antibodies (Biolegend) and rotated at 4°C for 4 hours prior to addition of protein G magnetic beads (Biorad, Hercules, CA) and overnight incubation at 4°C while rotating. Beads were magnetized and non-HA containing supernatants were removed prior to washing with a high salt buffer (50mM Tris, 300mM KCl, 12mM MgCl2, 10% NP-40, 1mM DTT, 100μ g/mL cycloheximide). Three washes were conducted prior to magnetizing the beads, removing all supernatants and then releasing HA bound transcripts from the beads using Buffer RLT supplemented with 2-β-mercaptoethanol from a Qiagen RNeasy microkit followed by in-column DNase I treatment and isolation of RNA according to manufacturer’s instructions. Samples were amplified for RNAseq by cDNA library preparation using a NuGen Ovation RNA v2 kit.

#### Illumina RNA sequencing and pathway analysis

A total of 16 mRNA samples were sequenced at Mayo Clinic Genome Facility using an Illumina HiSeq 4000. Reads were mapped to the mouse genome mm10 and reads per kilobase per million mapped reads (RPKM) were generated for each transcript. RPKM values were log2 transformed and low abundance transcripts were removed from further analysis (log2 RPKM less than -2 across all experimental groups were considered too lowly expressed). Differential expression, principal component analysis (PCA), and hierarchical clustering were performed using Partek Genomics Suite (Partek Inc., St. Louis, MO) with ANOVA and multiple comparison adjustment of p values using *Benjamini–Hochberg-Yekutieli* correction with false discovery rate (FDR) set at 5%. Pathway analyses were performed using DAVID (NIAID, Bethesda, MD) to examine significantly enriched pathways.

## Acknowledgements

Funding sources for JDF: Mayo Foundation, GHR Foundation, Mayo Clinic Center for Individualized Medicine, Mayo Clinic Gerstner Family Career Development Award, Ed and Ethel Moore Alzheimer’s Disease Research Program of Florida Department of Health (6AZ06), and NIH NS094137, AG047327, and AG049992. Funding sources for SK: The Robert and Clarice Smith and Abigail Van Buren Alzheimer’s Disease Research Program Fellowship, Mayo Clinic Program on Synaptic Biology and Memory, and NIH MH103632.

The authors declare no competing financial interests.

